# A salvaging strategy enables stable metabolite provisioning among free-living bacteria

**DOI:** 10.1101/2021.12.14.472564

**Authors:** Sebastian Gude, Gordon J. Pherribo, Michiko E. Taga

## Abstract

All organisms rely on complex metabolites such as amino acids, nucleotides, and cofactors for essential metabolic processes. Some microbes synthesize these fundamental ingredients of life *de novo*, while others rely on uptake to fulfill their metabolic needs. Although certain metabolic processes are inherently ‘leaky’, the mechanisms enabling stable metabolite provisioning among microbes in the absence of a host remain largely unclear. In particular, how can metabolite provisioning among free-living bacteria be maintained under the evolutionary pressure to economize resources? Salvaging, the process of ‘recycling and reusing’, can be a metabolically efficient route to obtain access to required resources. Here, we show experimentally how precursor salvaging in engineered *Escherichia coli* populations can lead to stable, long-term metabolite provisioning. We find that salvaged cobamides (vitamin B_12_ and related enzyme cofactors) are readily made available to non-productive population members, yet salvagers are strongly protected from overexploitation due to partial metabolite privatization. We also describe a previously unnoted benefit of precursor salvaging, namely the removal of the non-functional, proliferation-inhibiting precursor. As long as compatible precursors are present, any microbe possessing the terminal steps of a biosynthetic process can, in principle, forgo *de novo* biosynthesis in favor of salvaging. Consequently, precursor salvaging likely represents a potent, yet overlooked, alternative to *de novo* biosynthesis for the acquisition and provisioning of metabolites in free-living bacterial populations.

## Introduction

The general mechanisms enabling stable, long-term metabolite provisioning among free-living bacteria remain poorly understood[1–3]. While host-associated bacteria may count on a consistent supply of metabolites from their host, their free-living counterparts, particularly in unstructured environments, may not. Not only do the latter have no reliable access to metabolites provided by a host, but they also lack simple means for positive assortment[4], i.e., the ability of metabolite providers to preferentially supply metabolites to population members that benefit the provider, which is typically enabled by structured environments. In the absence of positive assortment, non-productive population members may easily invade, overexploit, and displace metabolite providers[5–8], causing an irreversible reduction in community diversity and metabolic capacity. The risk of overexploitation is not limited to fully public goods, but may even occur when providers possess the ability to retain exclusive access to some portion of the metabolite pool, as this private pool alone may be insufficient to support their long-term proliferation.

Bacteria require a diverse set of metabolites. These essential ingredients of life can be obtained through *de novo* biosynthesis or via uptake from the environment[9]. Metabolite provisioning among bacteria is frequently enabled by an intricate network of metabolic interactions[10]. Members of complex microbial communities can be broadly categorized according to their metabolic archetype for a particular metabolite as producers, degraders, salvagers, dependent consumers, and independents (Fig. 1A; this study focuses exclusively on the interplay between salvagers and dependent consumers). While independents, by definition, do not partake in interactions involving the metabolite, dependent consumers (i.e., non-productive population members) critically rely on metabolically active producers, degraders, or salvagers to fulfill their metabolic needs. Since certain biological functions are inherently ‘leaky’[11–14], some metabolites, such as degradation products of extracellularly processed polypeptides and polysaccharides[15, 16] or byproducts of overflow metabolism[17], are readily supplied as ‘public goods’. Yet, this type of metabolite provisioning is inherently limited to particular classes of metabolites, such as externally processed polymeric compounds.

**Figure 1:**
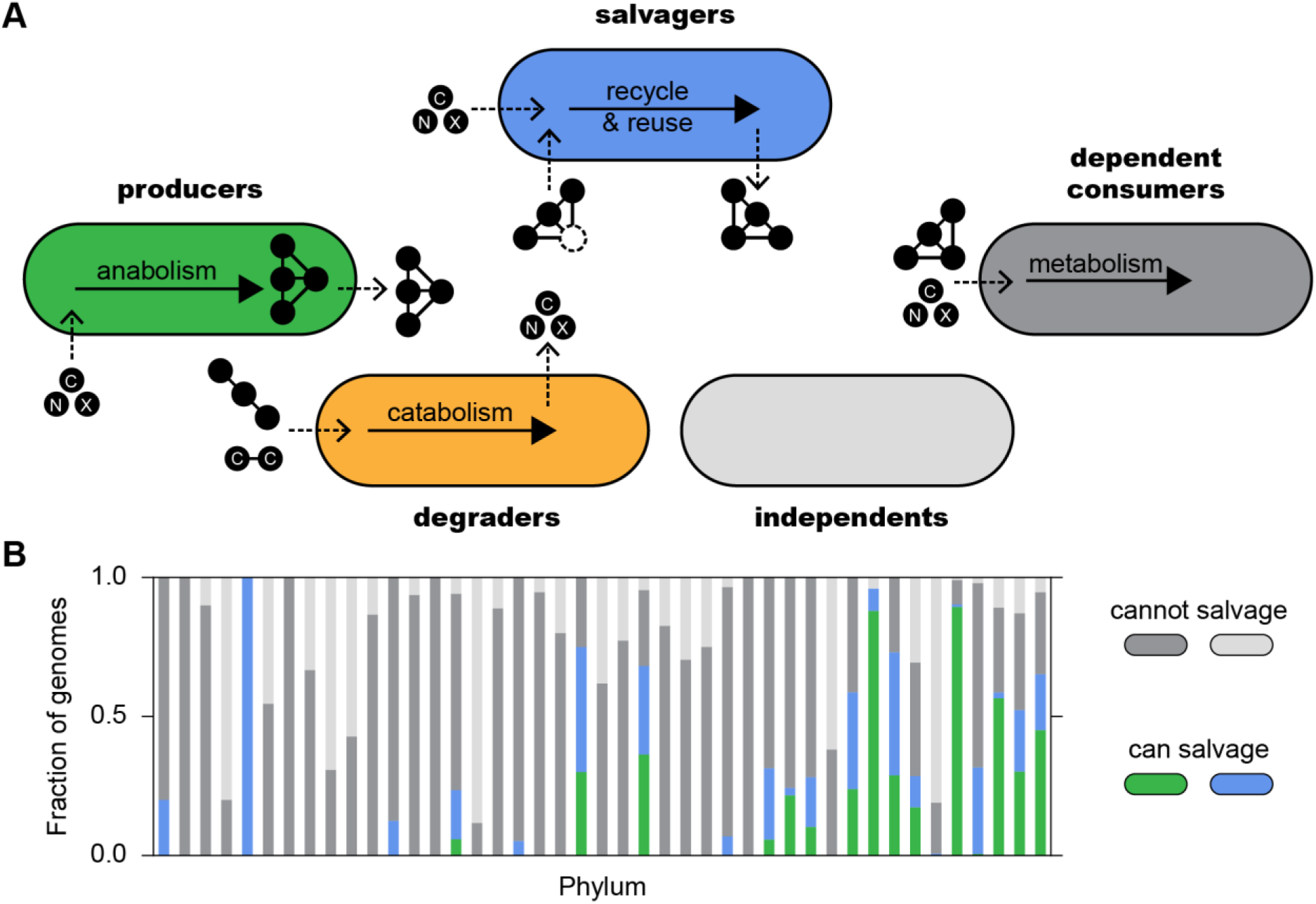
Metabolic archetypes in bacterial populations **A)** Illustration of distinct metabolic archetypes in bacterial populations. Producers create complex metabolites (black circles connected by lines) *de novo* from simple precursors (black circles) such as carbon- and nitrogen-containing compounds. Synthesized complex metabolites may be released and may benefit others. Dependent consumers take up simple compounds and complex metabolites to fulfill their metabolic needs. Dependence on complex metabolites may be facultative or obligate. Dependent consumers do not release metabolites. Degraders break down complex compounds such as polysaccharides or polypeptides. Degradation products are often accessible to others. Salvagers take up compounds from the environment and complete metabolite biosynthesis. Salvaged compounds may be released to the benefit of others. Independents neither create nor utilize a given metabolite. The metabolic archetype of an organism varies for different metabolites, e.g., a producer of one metabolite may be a consumer, degrader, salvager, or even be independent of another metabolite. **B.** Distribution of predicted cobamide archetypes among sequenced bacteria[30]. Colors as defined in A. Phyla are listed in Supplementary Table 1.

An alternative mode of gaining access to metabolites is salvaging, the process of ‘recycling and reusing’[18]. Bacterial salvaging is most commonly associated with iron acquisition[19–21]. In aerobic environments, many microbes release high-affinity iron-chelating siderophores to gain access to otherwise insoluble Fe^3+^. Yet, bacterial salvaging is by no means limited to the uptake and reuse of inorganic substances such as metals. Bacteria can also take up a variety of environmental metabolites in lieu of synthesizing them *de novo*. Classes of metabolites known to be salvaged include sugars, amino acids, nucleotides, and vitamins[22–27]. Notably, metabolite salvaging is not restricted to complete, terminal metabolites. Chemically stable precursors, for example intermediates of central metabolism or compounds involved in specialized metabolism, can be salvaged as well[28, 29].

While the genetic and biochemical basis of salvaging is well understood for many metabolites due to in-depth studies in a few bacterial species, less is known about the ecological ramifications of salvaging in multi-member bacterial populations[22–26, 28]. Here, as an example, we focus on the salvaging of the cobamide precursor, cobinamide (Cbi), and explore how the salvaging of metabolite precursors can shape bacterial population dynamics and stability. Cobamides (vitamin B_12_ and related enzyme cofactors) are cobalt-containing cofactors used in diverse metabolic pathways, and cobamide precursor salvaging is widely predicted among sequenced bacterial genomes[30] (Fig. 1B, Supp. Table 1). In this process, the incomplete and non-functional cobamide precursor Cbi is taken up and, in the presence of the required gene products for cobamide precursor salvaging, combined with a nucleoside containing a ‘lower ligand’ base of variable structure to form a complete, functional cobamide. By studying cobamide precursor salvaging, we demonstrate that salvaging can be an effective strategy enabling stable metabolite provisioning. Salvagers readily release assembled complete cobamides into the environment, where they can be utilized by non-productive population members, yet surprisingly we found that salvagers were strongly protected from overexploitation. These findings indicate that precursor salvaging may be a highly effective, generalizable way of enabling metabolite provisioning, even when means for positive assortment are absent.

## Results

### Salvaged metabolites are readily provisioned

To experimentally investigate precursor salvaging in the context of mixed free-living bacterial populations, we generated cobamide-dependent *Escherichia coli* populations (Fig. 2A, see Methods for details). Salvagers (Sal) only proliferated in glycerol minimal medium when either the complete cobamide vitamin B_12_ (B_12_) or the precursor Cbi and the lower ligand, 5,6-dimethylbenzimidazole (DMB), which together can be salvaged and assembled into functional B_12_ inside the cell, were supplied (Fig. 2B, blue symbols). As expected, no proliferation was observed in the presence of Cbi and DMB for dependent consumers (Dep) which have a disrupted cobamide precursor salvaging pathway (Fig. 2B, gray symbols).

**Figure 2:**
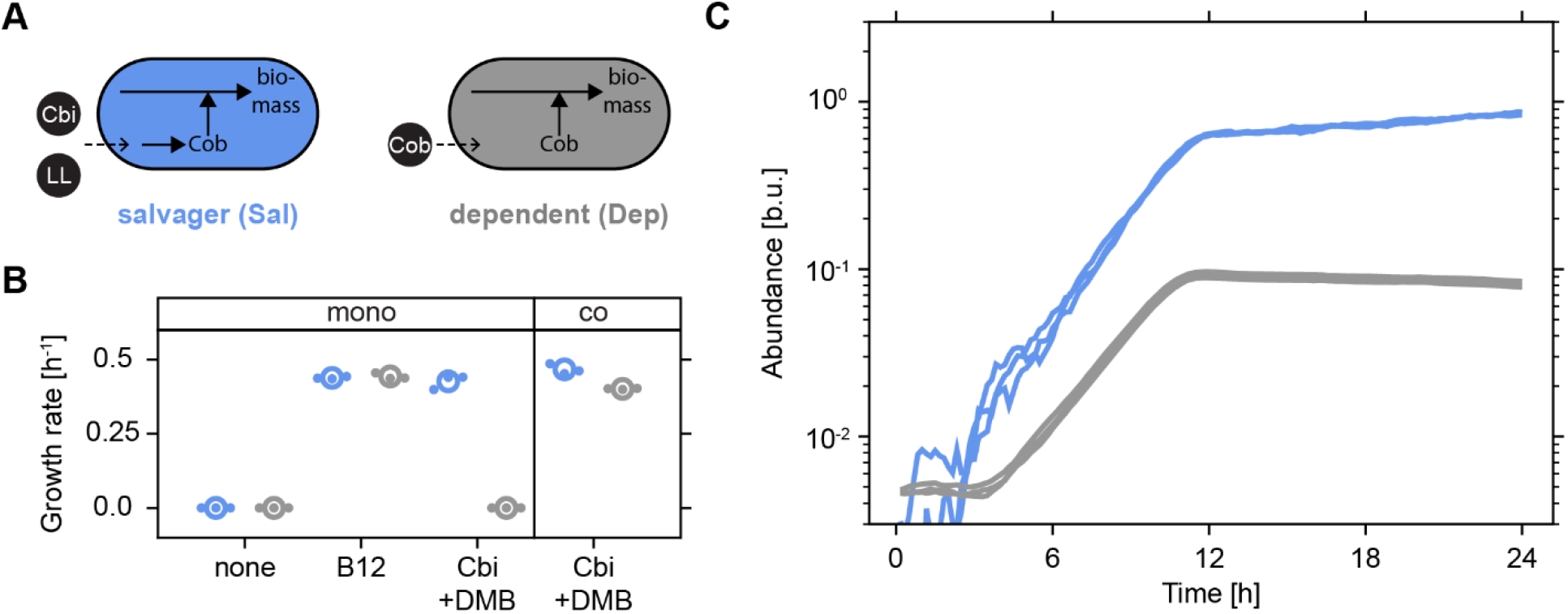
Salvaged cobamides are readily provisioned **A)** Illustration of cobamide archetypes. **B)** Exponential growth rates of salvagers (Sal, blue) and dependents (Dep, gray) in glycerol minimal medium with various additions (10 nM each). Strains were cultured alone (mono) or together (co, initial ratio: 1:1). Global means are shown as open circles and biological replicates as dots. N=3. **C)** Proliferation of jointly cultured salvagers (blue) and dependents (gray). Initial ratio: 1:1. N=3. Abbreviations: incomplete, non-functional cobamide precursor cobinamide (Cbi); lower ligand, a required component of a cobamide (LL); complete, functional cobamide (Cob); vitamin B_12_ (B12); 5,6-dimethylbenzimidazole, the lower ligand of B_12_ (DMB); bacterial units (b.u., see Methods for details).

Mixed salvager-dependent populations proliferated in the presence of Cbi and DMB (Fig. 2B). While salvagers were expected to proliferate, it remained unclear whether dependents would also be able to proliferate given that they rely on external provisioning of the complete cobamide B_12_. Previous studies reported a need to engineer or evolve ‘overproduction’ strains to enable provisioning of intracellular metabolites such as amino acids[11, 31]. Yet, conversely, a recent report demonstrated the general potency of vitamins and cofactors to promote stable metabolic interactions[32]. To dissect individual contributions to the overall increase in population size of the mixed salvager-dependent population, we established a calibrated fluorescence-based assay to track the dynamics of each subpopulation in well-stirred culture conditions over time (Supp. Fig. 1). The increase in the total population size was found to be driven by proliferation of both salvagers and dependents, indicating that the salvaged cobamides were readily released into the environment as a ‘public good’ and made available (i.e., provisioned) to the dependents (Fig. 2B and C).

Details of the proliferation dynamics markedly differed between the two subpopulations. Salvagers proliferated with no lag and experienced a growth rate of ~0.47 h^−1^ (Fig. 2B and C), comparable to their performance when cultured alone (Fig. 2B). Notably, dependents experienced an extended lag before proliferation commenced at a slightly reduced growth rate of ~0.40 h^−1^, and their final abundance, which they attained at the same time as salvagers ceased to proliferate, was only about 1/10 of the level achieved by salvagers (Fig. 2C). These findings suggested that the benefits of the salvaged complete cobamides were asymmetrically accessible to salvagers and dependents, even though mechanisms for positive assortment[4], such as spatial exclusion of dependents in structured environments[6], were inaccessible under these (well-stirred) culture conditions.

### Salvaging ‘preference’ determines population dynamics

We next explored the potential limits of the unprompted cobamide release and provisioning observed in mixed salvager-dependent populations. We reasoned that salvaging-based proliferation and metabolite provisioning should depend on the quantity and chemical identity of the available precursors. To test this hypothesis, we systematically varied external lower ligand supplementation and monitored proliferation of jointly cultured salvagers and dependents. Proliferation dose response curves in the presence of Cbi and various levels of DMB indicated that salvagers successfully proliferated at all tested concentrations of the externally supplied lower ligand (Fig. 3A). Conversely, proliferation of dependents ceased completely below 100 pM of externally supplied DMB, again pointing towards a preferential access of salvagers to the assembled cobamide. A similar trend was observed when a chemically different lower ligand, 2-methyladenine (2MA, lower ligand of the cobamide factor A, FA), was supplied (Fig. 3B). Once more only the proliferation of the dependents ceased at low levels of external lower ligand supplementation while salvagers continued to proliferate. Notably, a ~300-fold higher level of 2MA compared to DMB was required to enable dependent proliferation, indicating that chemical identity indeed played an important role. We reasoned that this difference in dependent proliferation between the two chemically distinct lower ligands could be caused by a difference in the metabolic ‘preference’ for the respective complete cobamides, which has been reported for several bacterial species[33–35]. Alternatively, a difference in the efficiency of the precursor salvaging pathway for installation of the two lower ligands could be decisive. Proliferation dose response curves of dependents cultured alone showed that a lower concentration of B12 compared to FA was required for dependent proliferation, yet this difference in metabolic ‘preference’ between the two complete cobamides was relatively minor (~2-fold) (Fig. 3C). This finding indicates that the drastic difference in dependent proliferation between the supplementation with the chemically distinct lower ligands DMB and 2MA was not due to a metabolic ‘preference’ for the complete cobamides B_12_ and FA, but was instead mainly caused by a difference in lower ligand ‘preference’ of the precursor salvaging pathway. Similar observations of a salvaging ‘preference’ in bacteria have previously been reported[36, 37].

**Figure 3:**
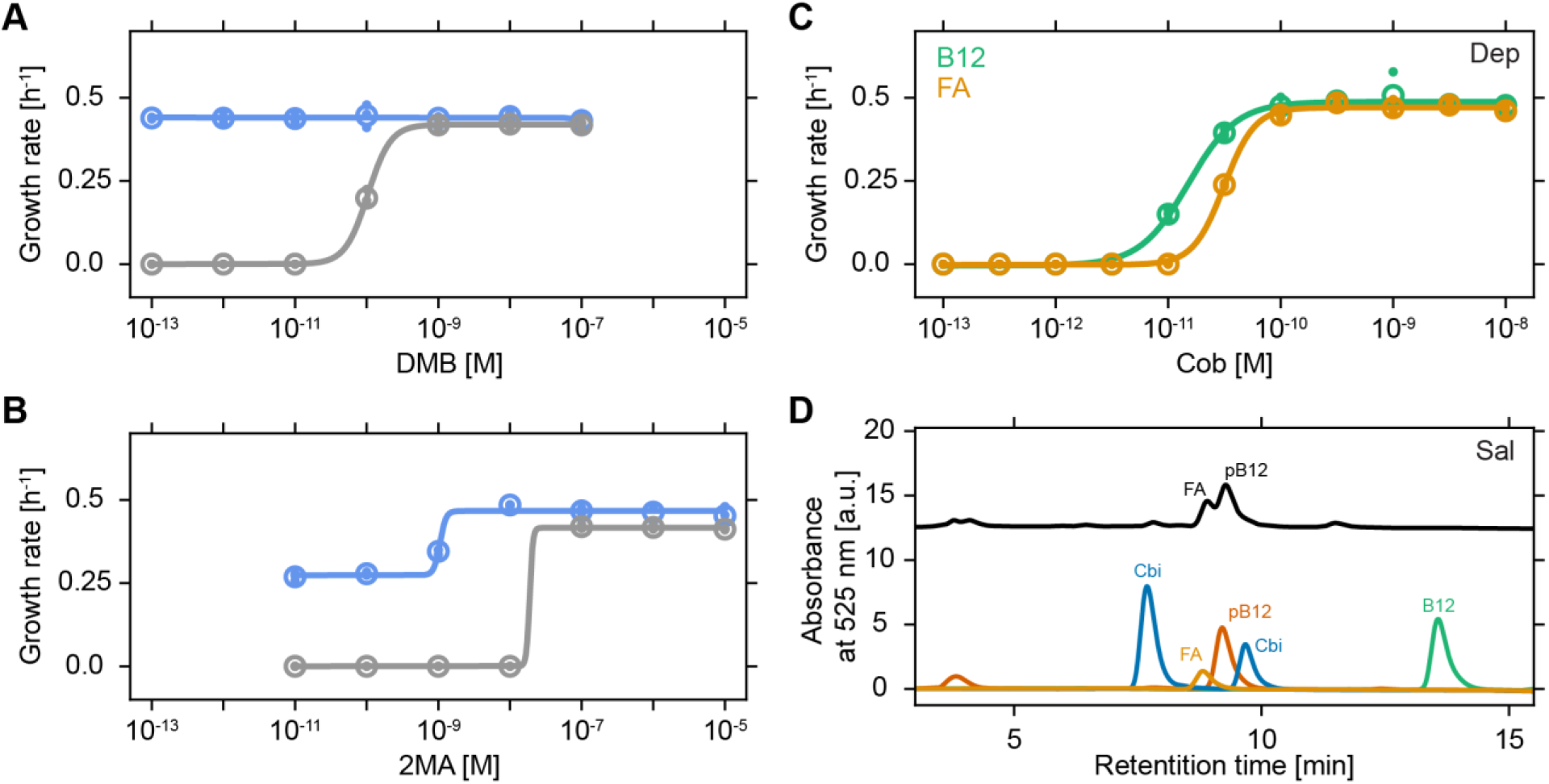
Salvaging ‘preference’ shapes population dynamics **A-B)** Growth rates of Sal (blue) and Dep (gray) when jointly cultured at an initial ratio of 1:1 in glycerol minimal medium supplemented with the cobamide precursor Cbi [10 nM] and varying concentrations of external lower ligand, DMB (A) and 2MA (B). **C)** Growth rate of Dep in glycerol minimal medium with varying concentrations of the complete cobamides B_12_ (green, EC_50_ = 1.5×10^−11^ M) and FA (yellow, EC_50_ = 3.1×10^−11^ M). EC_50_ is the cobamide concentration at half-maximal growth rate. In panels A-C, global means are shown as open circles, biological replicates as dots, and sigmoidal fits as lines. N=3. **D)** HPLC analysis of cell extracts of Sal cultured in glycerol minimal medium supplemented with Cbi [10 nM] in the absence of an externally supplied lower ligand (black). Standards of cobinamide (Cbi, blue), factor A (FA, yellow), pseudo-B_12_ (pB12, red), and vitamin B_12_ (B12, green) are shown. Abbreviations: a.u., arbitrary units.

The proliferation dose response curves of jointly cultured salvagers and dependents contained an additional noteworthy detail (Fig. 3A and B). Namely, proliferation of salvagers was observed for levels of externally supplied lower ligand that, even if conversion to complete cobamide was 100% efficient, were below the minimal level of externally supplied complete cobamide required for proliferation (Fig. 3C). Thus, salvagers were able to generate proliferation-supporting levels of complete cobamides in the absence of sufficient external lower ligand supplementation, presumably by incorporating internally produced lower ligands. Using high-performance liquid chromatography (HPLC) on cell extracts obtained from salvagers cultured in the presence of Cbi with no externally supplied lower ligand, we identified these ‘default’ cobamides as a mixture of FA (lower ligand: 2MA) and pseudo-B_12_ (pB_12_, lower ligand: Adenine) based on comparison of their respective retention times and characteristic peak spectra with purified standards (Fig. 3D, Supp. Fig. 4). Thus, as observed previously[38], salvagers can supplement limiting external precursor supply by tapping into their internal metabolite pool to obtain *de novo* synthesized lower ligands (here 2MA and adenine) to generate required complete cobamides.

### Salvaging can eliminate negative effects of metabolite accumulation

Precursor salvaging seemed not to incur a noticeable metabolic cost under our culture conditions, as salvagers proliferated at comparable growth rates in the presence of either the complete cobamide B_12_ or the precursors Cbi and DMB (Fig. 2B). Nevertheless, the fact that metabolically valuable salvaged complete cobamides were readily released and provisioned to dependents remained puzzling, particularly because it invites overexploitation by dependents which may cause a collapse of the entire mixed population[5–8]. In an attempt to gain some basic understanding of the factors that may facilitate the unprompted release of the metabolically valuable salvaged cobamides, we tried taking a more abstract perspective on the precursor salvaging process by comparing its functioning to other similarly structured cellular activities. On the most basic level, cobamide precursor salvagers take up metabolites, modify them intracellularly, and finally release them back into the extracellular environment. While the unprompted release of metabolically valuable salvaged metabolites begs an explanation, the similarly structured process of antibiotic inactivation certainly does not. Here, toxic compounds are taken up, internally converted into less harmful forms, and subsequently released into the environment[39]. We wondered whether in addition to its ‘constructive’ characteristic, i.e., the generation of complete, functional cobamides, cobamide precursor salvaging could also have benefits through its ‘consuming’ characteristic, i.e., the assimilation of the incomplete, non-functional cobamide precursor Cbi. The latter could be beneficial if the presence of high levels of metabolically valuable precursors interferes with proliferation, for example, if structural similarities between Cbi and B12 interfere with cellular functions via competitive binding to enzymes. Accumulation to proliferation-inhibiting levels has been noted for various metabolites[17, 40] and their consumption has been reported to stabilize commensal interactions[41].

Proliferation dose response curves of dependents cultured alone on various levels of the complete cobamide B_12_ in the absence and presence of high levels (10-fold higher than the highest tested level of B_12_) of Cbi indeed revealed a clear proliferation-inhibiting effect (Fig. 4A). The presence of high levels of environmental Cbi not only increased the EC_50_ (i.e., the concentration supporting half-maximal growth rate) of the proliferation dose response curve by about 5-fold (Fig. 4A, black arrow) but also induced the appearance of a drastic lag before proliferation was initiated for the lowest B_12_ levels that supported proliferation in the presence of surplus Cbi (Fig. 4B). Thus, though Cbi is metabolically beneficial as a precursor for salvaging, it clearly has negative effects when present in excess.

**Figure 4:**
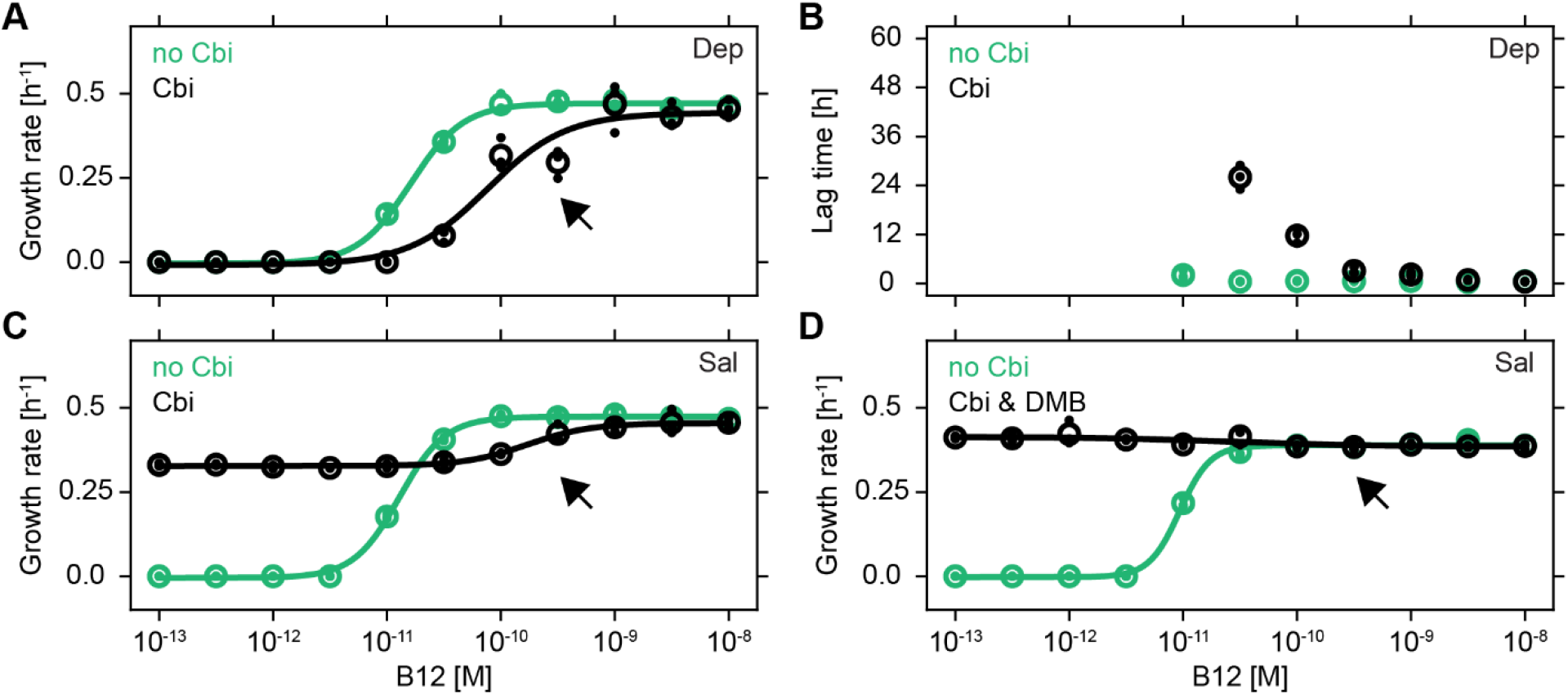
Salvaging can relieve proliferation inhibition **A-C)** Growth rate (A) and lag time (B) of Dep and growth rate of Sal (C) in glycerol minimal medium as a function of the concentration of B_12_ in the absence (green) and presence (black) of the cobamide precursor Cbi [10^−7^ M]. Global means are shown as open circles, biological replicates as dots, and sigmoidal fits as lines. N=3. **D)** Same conditions as C, but with the lower ligand DMB [10^−6^ M] also added (black). N=3.

We reasoned that this proliferation-inhibition caused by surplus Cbi could potentially be overcome, or at least be diminished, by the ‘consuming’ character of precursor salvaging. Indeed, salvagers cultured under the same conditions as described above did not exhibit the lag previously observed for dependents (Supp. Fig. 2, Fig. 4B). Yet, the inhibitory effect of Cbi seen at higher B_12_ levels remained unchanged (Fig. 4C, black arrow). Cobamide precursor salvaging in the absence of external lower ligand supplementation is likely limited by the internal availability of *de novo* synthesized lower ligands, which, under our conditions, could be insufficient to assimilate sufficient amounts of the proliferation-inhibiting Cbi. As a direct test, we supplied salvagers with a 10-fold excess of externally supplied lower ligand. Under these conditions salvagers were indeed able to completely relieve the detrimental effects of surplus Cbi (Fig. 4D). Consequently, the ‘consuming’ character of precursor salvaging, i.e., the uptake and assimilation of proliferation-inhibiting precursors and the subsequent export of complete cobamides, can be beneficial to salvagers, and thus may (partially) explain the unprompted release of salvaged metabolites.

To explore whether these Janus-like (i.e., two-sided) benefits of precursor salvaging were reflected in its regulatory logic, we constructed a fluorescent reporter for the expression of the major genes involved in the cobamide precursor salvaging process (i.e., cobUST) to determine whether salvaging is regulated by precursor abundance. While gene expression changed slightly (~4-fold) over time, we did not observe differential gene expression between conditions that required precursor salvaging (supplementation of Cbi and DMB) and conditions that did not (supplementation of B_12_) (Supp. Fig. 3). Thus, our results suggested that cobamide precursor salvaging in *E. coli* was not regulated at the level of gene expression under our culturing conditions, and that the precursor salvaging pathway was expressed independently of the need for cobamide precursor salvaging.

### Partial privatization enables salvagers to persist stably

Unprompted metabolite release in the absence of mechanisms for positive assortment bears the risk of overexploitation, potentially causing an associated collapse of a metabolically interconnected population if non-productive members (i.e., dependents) possess a proliferation advantage over their productive counterparts (i.e., salvagers)[42]. In particular, faster proliferating dependents could cause an arrest in salvager proliferation, which would inevitably result in a collapse of the entire metabolically interconnected population, by quickly depleting the publicly available cobamide. Alternatively, salvagers could be able to persist independently of the dependents’ proliferation properties by securing an adequate access to cobamides prior to releasing them into the extracellular environment. As an initial step to assess the risk of population collapse in mixed salvager-dependent populations, we created faster-proliferating mutant variants by moving the salvager and dependent genotypes into a closely related *E. coli* MG1655 GB-1 mutant background (Sal^GB-1^ and Dep^GB-1^)[43]. The GB-1 background was contains only two mutations (in the glycerol kinase *glpK* and the RNA polymerase β’ *rpoC*) relative to wild type *E. coli* MG1655 and confers a growth rate advantage of ~45% compared to the original salvager and dependent in glycerol minimal medium (compare Figs. 5A and 2B)[43]. Combinatorial culturing of all four possible salvager-dependent pairs as 1:1 mixtures resulted in expected changes in the final population composition (Fig. 5B). Specifically, when jointly cultured with Sal, Dep only reached about 1/10 of the abundance of Sal after 24 hours, in agreement with our previous observations (Fig. 2C). Jointly cultured Sal^GB-1^ and Dep^GB-1^ showed a similar pattern, indicating that only relative, and not absolute, proliferation characteristics determined the population composition. Dep proliferation was much reduced when it was cultured with the faster proliferating Sal^GB-1^ mutant, whereas the contrary effect was observed if the dependent (Dep^GB-1^) had a proliferation advantage over the salvager (Sal). Together, these findings demonstrated how relative changes in the maximally achievable growth rate can drastically reshape the composition of metabolically linked populations. Moreover, it suggested that dependents may indeed have the ability to overexploit salvagers by quickly depleting the public cobamide pool if they have sufficiently large growth rate advantages.

**Figure 5:**
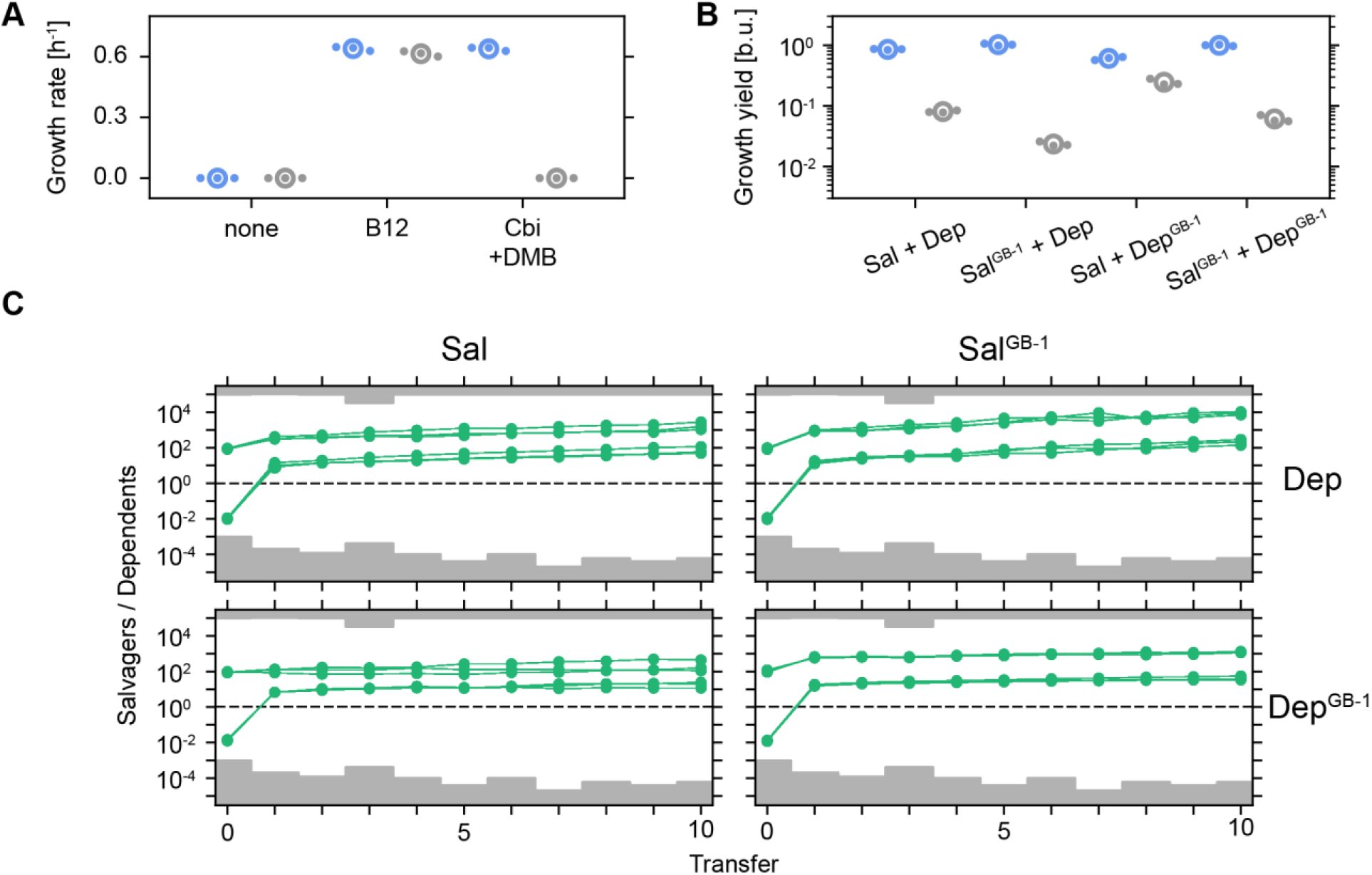
Partial cobamide privatization protects salvagers from overexploitation **A)** Growth rates of Sal^GB-1^ (blue) and Dep^GB-1^ (gray) when cultured alone in glycerol minimal medium with various additions (10 nM each). **B)** Growth yields after 24 hours of jointly cultured salvagers (Sal or Sal^GB-1^, blue) and dependents (Dep or Dep^GB-1^, gray) in glycerol minimal medium supplemented with Cbi [10 nM] and DMB [10 nM]. Initial ratio: 1:1. In panels A and B, global means are shown as open circles and biological replicates as dots. N=3. **C)** Population ratio (green dots) as function of transfer for jointly cultured salvagers (Sal or Sal^GB-1^) and dependents (Dep or Dep^GB-1^) undergoing daily 1:100 [v:v] proliferation-dilution cycles in glycerol minimal medium supplemented with Cbi [10 nM] and DMB [10 nM]. Solid green lines connect individual cultures through subsequent transfers. Gray areas mark transfer-specific estimates of population ratio detection limits (for details see Methods). N=3.

To explore the consequences of growth rate differences on long-term population dynamics and stability more rigorously, we exposed all four mixed salvager-dependent populations to a sequence of proliferation-dilution cycles (Fig. 5C). Mixed populations were initialized with either salvagers or dependents 100-fold in the majority, with a fixed total initial population size. The culture medium including glycerol, Cbi, and DMB was replenished at each transfer. Interestingly, salvagers were able to persist, and even achieved clear numerical dominance, in all four mixed populations (Fig. 5C). These findings demonstrated that salvagers were strongly protected from overexploitation even if dependents were able to proliferate faster by as much as 45%. Details of long-term population dynamics varied between the four mixed populations. Differences in maximally achievable growth rate was a dominant factor determining the rate at which population ratios changed over time. Faster-proliferating salvagers outcompeted slower-proliferating dependents at a higher rate (Fig. 5C, Sal^GB-1^/Dep) compared to slower-proliferating salvagers (Fig. 5C, Sal/Dep and Sal/Dep^GB-1^). Similarly, faster-proliferating dependents fared better than their slower-proliferating counterparts, as they were outcompeted at a lower rate (Fig. 5C). Interestingly, the population ratio (from transfer 1 onwards) of Sal/Dep changed at a slightly higher rate than that of Sal^GB-1^/Dep^GB-1^, despite the observation that there was no difference in maximally achievable growth rate within each of the strain pairs (Fig. 2B, Fig. 5A and C). This finding may indicate that, counter to a commonly held assumption and our findings in single batch cultures (Fig. 5B), long-term competition dynamics in this metabolically interconnected microbial system were not solely determined by relative differences in growth rate but rather absolute growth rates may play a role as well. Alternatively, the observed difference in long-term competition dynamics could also be explained by phenotypic variations other than differences in exponential growth rate in the strain backgrounds. For example, differences in cobamide release during stationary phase could shift the long-term population dynamics to the benefit of the dependents. The faster proliferating GB-1 mutant was originally obtained through controlled laboratory evolution experiments in which it was constantly kept in exponential phase[43]. Detailed multi-omics characterization of the GB-1 mutant indeed indicated that its proteome shows a clear reduction in stationary phase and stress-induced proteins[44]. The latter may cause a reduction in survival probability when populations, as in our experimental setting and likely in many natural settings, spent a considerable time in stationary phase. A repeated occurrence of substantial cell death, together with its associated release of intracellular material, may enable the dependents to obtain access to otherwise sequestered metabolites due to the disintegration of cells during stationary phase.

Together, these findings indicate how intracellular metabolic processes that innately enable (partial) retention of metabolites inside the cell are fundamentally distinct from extracellular processes like degradation of some polysaccharides[15]. In the latter case, metabolites immediately become ‘public goods’ that have to be taken up from the environment, thus inherently limiting the ability to privatize resources due to the finite affinities of import systems[45, 46]. In contrast, intracellular metabolic processes, such cobamide precursor salvaging, may initialize metabolite ‘leakage’ only once the internal metabolite level reaches a certain threshold, thus ensuring preferential access due to partial metabolite privatization. Ultimately, our findings demonstrate the ability of intracellular metabolic processes to establish stable metabolic interactions, with important ramifications on community stability and the protection of productive population members.

## Discussion

Cobamide precursor salvaging has been experimentally observed and is predicted to be widespread among bacteria[30, 47]. Here, we showed that (1) salvaging of cobamide precursors does not incur a noticeable metabolic burden, (2) salvaged cobamides are readily released and provisioned to others, (3) cobamide precursor salvaging provides benefits through the generation of complete functional cofactors as well as through the removal of non-functional, proliferation-inhibiting precursors, and (4) organisms engaging in cobamide precursor salvaging are intrinsically protected from overexploitation due to partial metabolite privatization. While likely not all of these features are transferable to salvaging of other types of metabolites, some certainly will. For example, removal of proliferation-inhibiting metabolites is likely widespread as high levels of many compounds, such as cysteine[40], can lead to disturbances of optimal metabolic fluxes and thus hinder proliferation. Furthermore, (partial) privatization, which has also recently been proposed in the context of other microbial activities[7, 48, 49], is likely applicable to a broad set of metabolites, including amino acids, nucleotides, and cofactors, that are inherently processed inside the cell. Thus, organisms generating these intracellular metabolites, either via salvaging or *de novo* biosynthesis, are likely naturally protected from overexploitation by non-productive population members, and therefore may facilitate the emergence and stable maintenance of metabolite provisioning interactions.

Salvaging, the concept of ‘recycling and reusing’, provides a resource-efficient way to use environmental metabolites and inorganic substances that were released due to metabolite overflow or cell lysis. It is important to note that metabolite salvaging is not limited to the uptake and completion of precursors, which was the focus of this study. Salvaging can also be combined with partial catabolism and metabolite modification to interconvert complete, yet enzymatically unfavorable, metabolites into more useful forms[50, 51]. Taken together, salvagers of various metabolites likely play an important, yet underappreciated, role in many natural bacterial populations by acting as hubs facilitating efficient community functioning by rerouting and interconverting environmental metabolic fluxes. Innate protection from overexploitation by non-productive population members positions salvaging of intracellular metabolites as an effective mechanism to act in a broad range of environmental conditions, even those in which positive assortment is not accessible (e.g., free-living populations).

## Acknowledgement

This work was funded by National Institutes of Health grants R01GM114535 and R35GM139633 to M.E.T. We thank O. Sokolovskaya for initially suggesting to employ *E. coli* mutant strains. We thank K. Mok and O. Sokolovskaya for assistance with HPLC measurements and H. Nolla at the Flow Cytometry Facility in the Cancer Research Laboratory at UC Berkeley for assistance with flow cytometry measurements. We thank members of the Taga lab for helpful discussions. We also thank K. Kennedy, K. Mok, and Z. Hallberg for critical reading of the manuscript. *E. coli* strain MG1655 GB-1 was a kind gift of B.O. Palsson.

## Author contributions

S.G. and M.E.T. conceived research. S.G., G.J.P., and M.E.T. developed *E. coli*-cobamide model system. S.G. designed and performed all experiments except corrinoid extraction and analysis. G.J.P. designed and performed corrinoid extraction and analysis. S.G. analyzed all data.

S.G. and M.E.T. interpreted the data and wrote the manuscript with input from G.J.P.

## Declaration of interests

The authors declare no competing interests.

## Resource availability

### Contact

Further information and requests for resources and reagents should be directed to and will be fulfilled by Michiko E. Taga.

### Materials availability

All materials generated in this study will be made available on request, but we may require a payment and/or a completed materials transfer agreement if there is potential for commercial application.

### Data and code availability

All data is available from the lead author upon reasonable request. All original code can be found in the Supplementary Information.

## Methods

### Strains and plasmids

All strains are derivatives of *E. coli* strain MG1655. Sal^GB-1^ and Dep^GB-1^ were derived from *Escherichia coli* MG1655 mutant GB-1[43]. Strain Sal^GB-1^ was constructed by introducing into strain GB-1 the *metE*::Kan^R^ allele from donor strain JW3805-1[53] via P1 transduction and subsequently removing the Kan^R^ marker by transient introduction of the plasmid pCP20 encoding the FLP recombinase[54]. To generate strain Dep^GB-1^, the *cobUST* operon was deleted via lambda red recombineering[54] and the gene *cobC* was replaced via P1 transduction as described above. After each step, the selective marker was removed. Gene deletions were verified via PCR and phenotyping of growth in glycerol minimal medium supplemented with either methionine, vitamin B_12_, cobinamide and DMB, or no addition.

Fluorescent marker and reporter plasmids are based on pETMini[51] and were assembled via isothermal cloning[55]. Insert sequences were verified via Sanger sequencing.

All bacterial strains and plasmids are listed in Supplementary Table 2 and 3.

### Media and culturing

For proliferation and serial passaging assays, 2 mL glycerol minimal medium (50 mM KPO_4_, 67 mM NaCl, 7.6 mM (NH_4_)_2_SO_4_, 500 μM MgSO_4_, 1.25 μM Fe_2_(SO_4_)_3_, 0.2% [v:v] glycerol, pH 7.4) supplemented with 1 g/L methionine was inoculated with single colonies from LB plates (10 g/L tryptone, 5 g/L yeast extract, 5 g/L NaCl, 15 g/L Bacto agar), and incubated at 37 °C with aeration (200 rpm) until saturation was reached after approx. 24 hours. Cultures were transferred into 2 mL fresh glycerol minimal medium supplemented with methionine as 1:100 [v:v] dilutions, incubated at 37 °C with aeration, and harvested in mid-exponential phase (approx. OD_600_ 0.3-0.6). Cells were washed three times by centrifugation at 14000 rpm followed by resuspension in unsupplemented minimal medium. Optical density (OD_600_) was adjusted to 10^−1^ and, if required, cultures were mixed 1:1 [v:v], or as indicated, to obtain mixed populations. If necessary, media were supplemented with 25 mg/L kanamycin to retain plasmids.

For proliferation assays, precultured and OD-adjusted cultures were transferred as 1:10 [v:v] dilutions into glass-bottom 96-well culture plates (CellVis) containing a final volume of 200 μL fresh glycerol minimal medium supplemented with either 10 nM vitamin B_12_, 10 nM cobinamide and 10 nM 5,6-dimethylbenzimidazole (DMB), or no additions, if not indicated otherwise. 96-well culture plates were sealed with an evaporation seal (Breathe-Easy® sealing membrane) and incubated at 37 °C with continuous shaking in a multi-well plate reader (Tecan Spark) for 36 or 60 hours. Absorbance at 600 nm and fluorescence (CFP: excitation 455/5 nm, emission 475/5 nm; YFP: excitation 514/5 nm, emission 550/30 nm) were recorded every 10 minutes. Fluorescence measurements were converted into units equivalent to OD_600_ (see below and Supplementary Figure 1). Fluorescence was recorded only for experiments containing two strains. All proliferation experiments were performed in biological triplicates started on the same day. Proliferation phenotypes showed little day-to-day variation.

For serial passaging assays, precultured and OD-adjusted mixed cultures were transferred as 1:10 [v:v] dilutions into clear plastic 96-well culture plates (Corning®) containing a final volume of 200 μL fresh glycerol minimal medium supplemented with 10 nM cobinamide and 10 nM DMB. Wells without cultures were filled with 200 μL dH2O. 96-well culture plates were sealed with an evaporation seal (AeraSeal^TM^, EXCEL Scientific) and incubated at 37 °C while shaking at 1200 rpm in a heated plate shaker (Southwest Science). Cultures were transferred every 24 hours as 1:100 [v:v] dilutions into fresh glycerol minimal medium supplemented with 10 nM cobinamide and 10 nM DMB. At each transfer samples were taken for composition analysis at the Flow Cytometry Facility in the Cancer Research Laboratory at UC Berkeley. Prior to flow cytometry analysis (BD LSRFortessa™ Cell Analyzer), samples were diluted 1:1000 [v:v] into unsupplemented minimal medium. Forward scatter (FSC; V=350, log), side scatter (SSC; V=280, log), FITC (V=650, log), and AmCyan (V=750, log) were recorded. Event detection was triggered by a threshold in SSC of 500. Samples were run at approx. 300-600 events per second, 10^5^ events were collected per sample, and lines were flushed with 10% bleach, rinse solution, and dH2O between sample runs. The serial transfer experiment was performed in biological triplicate started on the same day.

For corrinoid extractions, 2 mL glycerol minimal medium supplemented with 100 mg/L methionine was inoculated with single colonies of strains grown on LB plates and incubated at 37 °C with aeration until saturation was reached. Cells were washed twice by centrifugation at 14000 rpm and resuspension in unsupplemented minimal medium, diluted 1:500 [v:v] into 500 mL glycerol minimal medium supplemented with 10 nM cobinamide, incubated at 37 °C with aeration for 24 hours, and harvested for metabolite extraction, purification, and profiling (see below).

### Evaluation of proliferation dynamics

For proliferation assays, population dynamics were evaluated in Python[56] (version 3.8) with the following steps. (1) Fluorescence-to-absorbance conversion (if applicable): Fluorescence-to-absorbance conversion factors were estimated in a day-strain-medium-dependent manner for each fluorescent channel. Conversion factors were obtained as replicate-averaged slopes of linear fits (*NumPy[57]* (version 1.21.2) function *polyfit*) of raw fluorescence vs raw absorbance (A_600_) in the range A_600_ > 0 and A_600_ < 0.3 of cultures containing only a single marker plasmid. Raw fluorescence reads were then converted into equivalent units of raw absorbance by multiplication of the conversion factors with the raw fluorescence reads. (2) Absorbance-to-abundance conversion: All reads were pathlength-corrected to obtain abundances in arbitrary bacterial units (b.u.; 1 b.u. is equivalent to an OD_600_ of 1). (3) Blank subtraction: For reads based on absorbance measurements, blanks were estimated in a well-specific manner as the difference of the mean of the first two time points and the known initial abundance. For reads based on fluorescence measurements, blanks were estimated in a global manner as means over all wells containing cultures that did not carry the specific fluorescent marker plasmid. (4) Smoothing and shortening: Blank-corrected abundance timeseries were smoothed with a centered rolling mean filter (*pandas* (version 1.3.2) *DataFrame* function *rolling*) using a window width of 7 data points with all data points equally weighted and a minimal window width of 1 point. Thereafter, smoothed abundance timeseries were shortened to 24 or 60 hours. (5) Extraction of proliferation parameters: Growth rate estimates were obtained via a Theil-Sen estimator (*SciPy[58]* (version 1.7.1) module *stats.mstats* function *theilslopes*) which was applied to log-transformed abundances. Estimation was restricted to data points for which the abundance was larger than 1.5×10^−2^ b.u. and smaller than 6×10^−2^ b.u. (Fig. 3C and Fig. 4ACD) or the abundance was larger than 1.5×10^−2^ b.u. and latest data point for which the rolling-window growth rate estimate (see below) was larger than 0.1 h^−1^ (Fig. 2B, Fig. 3AB, and Fig. 5A). Growth rate was set to zero if fewer than 7 data points fell into the estimation interval. Growth rate estimates were generally found to be reproducible within +/− ~0.025 h^−1^ (day-to-day and channel-to-channel variability). Rolling-window growth rate estimates were obtained by applying a Theil-Sen estimator (*SciPy[58]* (version 1.7.1) module *stats.mstats* function *theilslopes*) to log-transformed abundances in a centered rolling window fashion (*pandas* (version 1.3.2) *DataFrame* function *rolling*) using a window width of 19 data points with all data points equally weighted and a minimal window width of 10 data points. Lag time estimates were obtained by linear interpolation of growth rates and intercepts obtained from the Theil-Sen estimator to known log-transformed initial abundances. Lag time was set to negative infinity for non-growing populations. (6) Extraction of dose response curve parameters (if applicable): Dose response curve parameters were obtained via non-linear fitting (*SciPy[58]* (version 1.7.1) module *optimize* function *curve_fit*) of extracted proliferation parameters to a four-parameter dose response equation:

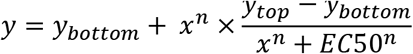

with variables *x* (effector concentration) and *y* (proliferation parameter), and parameters *y*_*bottom*_ (proliferation parameter value for *x* → 0), *y*_*top*_ (proliferation parameter value for *x* → *∞*), *n* (Hill coefficient), and *EC*50 (half maximal effective concentration).

### Evaluation of proliferation-dilution cycles

Flow cytometry data was analyzed in custom Python[56] (version 3.8) scripts using the FlowCal[59] (version 1.3.0) package. Briefly, data were loaded and transformed into arbitrary units via FlowCal. Gating was directly performed in Python. Events were classified as YFP-positive-CFP-negative if FITC > 1×10^3^ a.u. and AmCyan < 4×10^3^ a.u., as YFP-negative-CFP-positive if FITC < 7×10^2^ a.u. and AmCyan > 2×10^3^ a.u., as YFP-negative-CFP-negative if FITC < 1×10^3^ a.u. and AmCyan < 2×10^3^ a.u., and as YFP-positive-CFP-positive otherwise. Some gating conditions were altered for transfer zero as cells pre-cultured in the presence of methionine and harvested in mid-exponential phase showed slightly different fluorescence properties than cells cultured in the presence of cobinamide and DMB and harvested in stationary phase. In particular, YFP-negative-CFP-positive if FITC < 7×10^2^ a.u. and AmCyan > 5×10^3^ a.u. and YFP-negative-CFP-negative if FITC < 1×10^3^ a.u. and AmCyan < 5×10^3^ a.u. All other gating was as described above. Population ratios (Sal: Dep) were determined as the ratio of the number of YFP-negative-CFP-positive events over the number of YFP-positive-CFP-negative events. The dynamic range of the population ratio estimation was determined in a transfer-specific manner by the ratio of the number of YFP-negative-CFP-positive events obtained when measuring a blank (or 1, if no such events where detected) over the total number of events collected per mixed cultures (i.e., 10^5^) and the ratio of the total number of events collected per mixed cultures (i.e., 10^5^) over the number of YFP-positive-CFP-negative events obtained when measuring a blank (or 1, if no such events where detected), respectively. For a discussion of YFP-negative-CFP-negative events and YFP-positive-CFP-positive events see Supplementary Figure 5.

### Corrinoid extraction and analysis

Corrinoid extraction, cyanation, and purification were performed as previously described[47, 60]. Briefly, 500 mL of cultures were harvested by centrifugation and corrinoids were extracted from the pellet with methanol and cyanated with 20 mg potassium cyanide per gram (wet weight) of cells. Following cyanation, samples were desalted using C18 SepPak (Waters Associates, Milford, MA). HPLC purification and profiling were performed as previously described[47].

## Supplemental information

### Scripts_Fig01B.zip

Data analysis scripts for data shown in Fig 1 B.

### Scripts_Fig02B_Cocultures.zip

Data analysis scripts for coculture data shown in Fig 2 B.

### Scripts_Fig02B_GrowtwhRate_Plotting.zip

Data analysis scripts for growth ate data shown in Fig 2 B.

### Scripts_Fig02B_Monocultures.zip

Data analysis scripts for monoculture data shown in Fig 2 B.

### Scripts_Fig2C.zip

Data analysis scripts for data shown in Fig 2 C.

### Scripts_Fig03A.zip

Data analysis scripts for data shown in Fig 3 A.

### Scripts_Fig03B.zip

Data analysis scripts for data shown in Fig 3 B.

### Scripts_Fig03C.zip

Data analysis scripts for data shown in Fig 3 C.

### Scripts_Fig03D.zip

Data analysis scripts for data shown in Fig 3 D.

### Scripts_Fig04AB.zip

Data analysis scripts for data shown in Fig 4 A and B.

### Scripts_Fig04C.zip

Data analysis scripts for data shown in Fig 4 C.

### Scripts_Fig04D.zip

Data analysis scripts for data shown in Fig 4 D.

### Scripts_Fig05A.zip

Data analysis scripts for data shown in Fig 5 A.

### Scripts_Fig05B.zip

Data analysis scripts for data shown in Fig 5 B.

### Scripts_Fig05C.zip

Data analysis scripts for data shown in Fig 5 C.

### Scripts_SuppFig01.zip

Data analysis scripts for data shown in Supp Fig 1.

### Scripts_SuppFig02A.zip

Data analysis scripts for data shown in Supp Fig 2 A.

### Scripts_SuppFig02B.zip

Data analysis scripts for data shown in Supp Fig 2 B.

### Scripts_SuppFig03.zip

Data analysis scripts for data shown in Supp Fig 3.

### Scripts_SuppFig04.zip

Data analysis scripts for data shown in Supp Fig 4.

### Scripts_SuppFig05.zip

Data analysis scripts for data shown in Supp Fig 5.

---

**Supplementary Figure 1:**
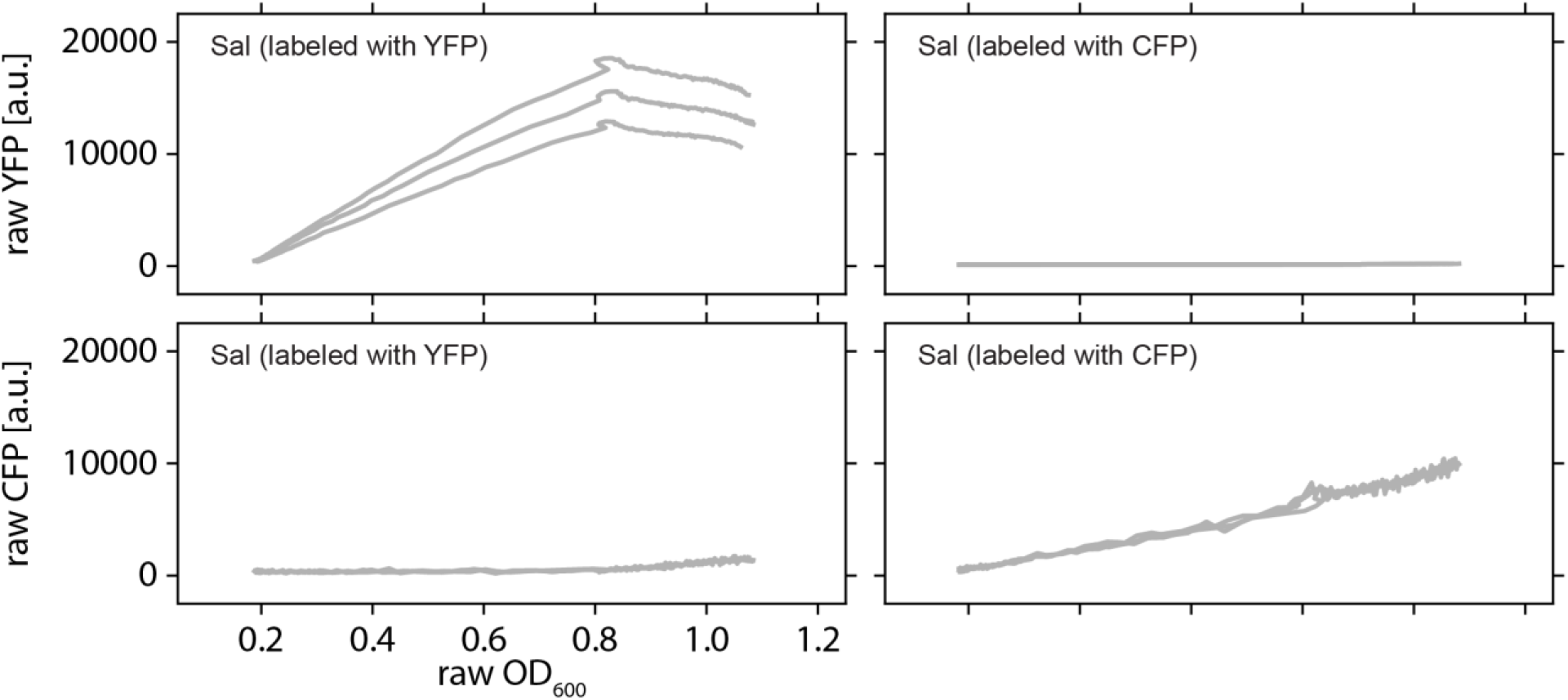
Calibrated fluorescence as proxy for abundance in mixed populations Examples of raw YFP and CFP fluorescence as function of raw optical density at 600 nm (OD_600_) extracted from proliferating, fluorescently labeled salvager populations (Sal) cultured in glycerol minimal medium supplemented with Cbi [10 nM] and DMB [10 nM]. Non-specific fluorescence background and crosstalk between fluorescence channels were small (bottom left and top right). YFP fluorescence underestimated abundances for raw OD_600_ >0.8. Day- and medium-specific fluorescence-to-abundance conversion factors were extracted as slopes from linear fits in the range 0 ≤ raw OD_600_ ≤ 0.6 and averaged over all biological replicates. The equivalent of a blank-corrected OD_600_ of 1 was defined as 1 b.u. (arbitrary bacterial unit). N=3. Abbreviations: arbitrary units (a.u.).

**Supplementary Figure 2:**
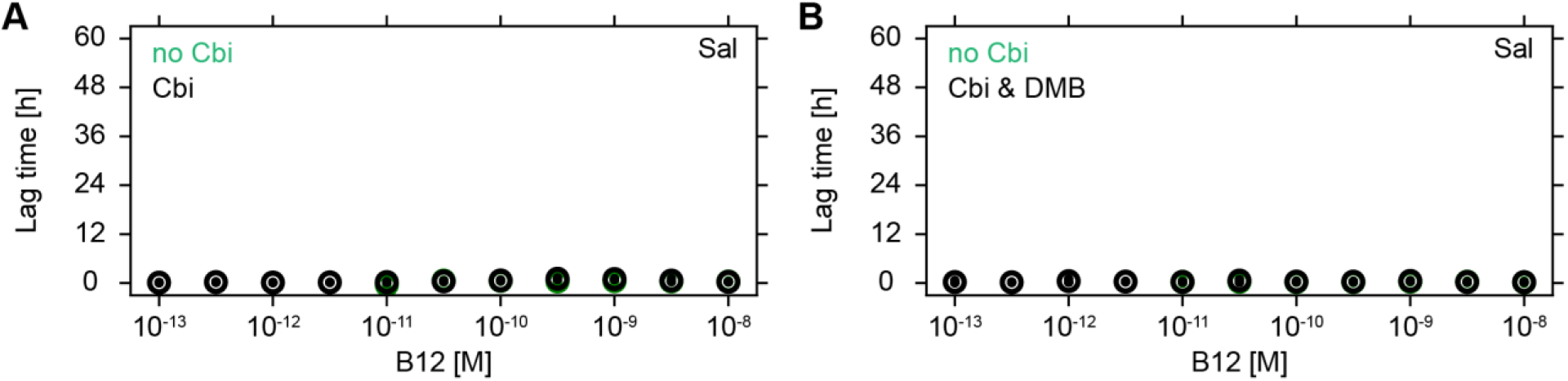
Salvagers do not exhibit lag **A)** Lag time of Sal in glycerol minimal medium as function of B_12_ concentration in presence (black) and absence (green, hidden behind black symbols) of Cbi [10^−7^ M]. **B)** Lag time of Sal in glycerol minimal medium as function of B_12_ concentration in presence (black) and absence (green, hidden behind black symbols) of Cbi [10^−7^ M] and DMB [10^−6^ M]. In panels A and B, global means are shown as open circles and biological replicates as dots. N=3.

**Supplementary Figure 3:**
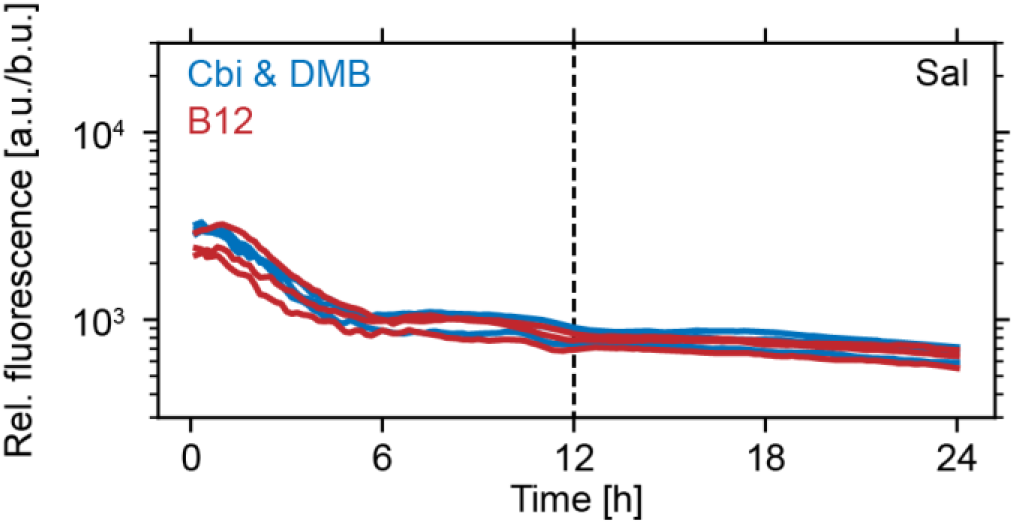
Cobamide precursor salvaging is not differentially regulated at the level of transcription Relative fluorescence (fluorescence in arbitrary units per bacterial abundance) of Sal carrying a YFP transcriptional reporter for cobUST (i.e., major cobamide salvaging genes) activity as function of time in glycerol minimal medium supplemented with Cbi [10 nM] and DMB [10 nM] (blue) or B_12_ [10 nM] (red). Dashed vertical line marks approximate time point at which proliferation ceases. N=3. Abbreviations: arbitrary units (a.u.) and bacterial units (b.u., see Methods for details).

**Supplementary Figure 4:**
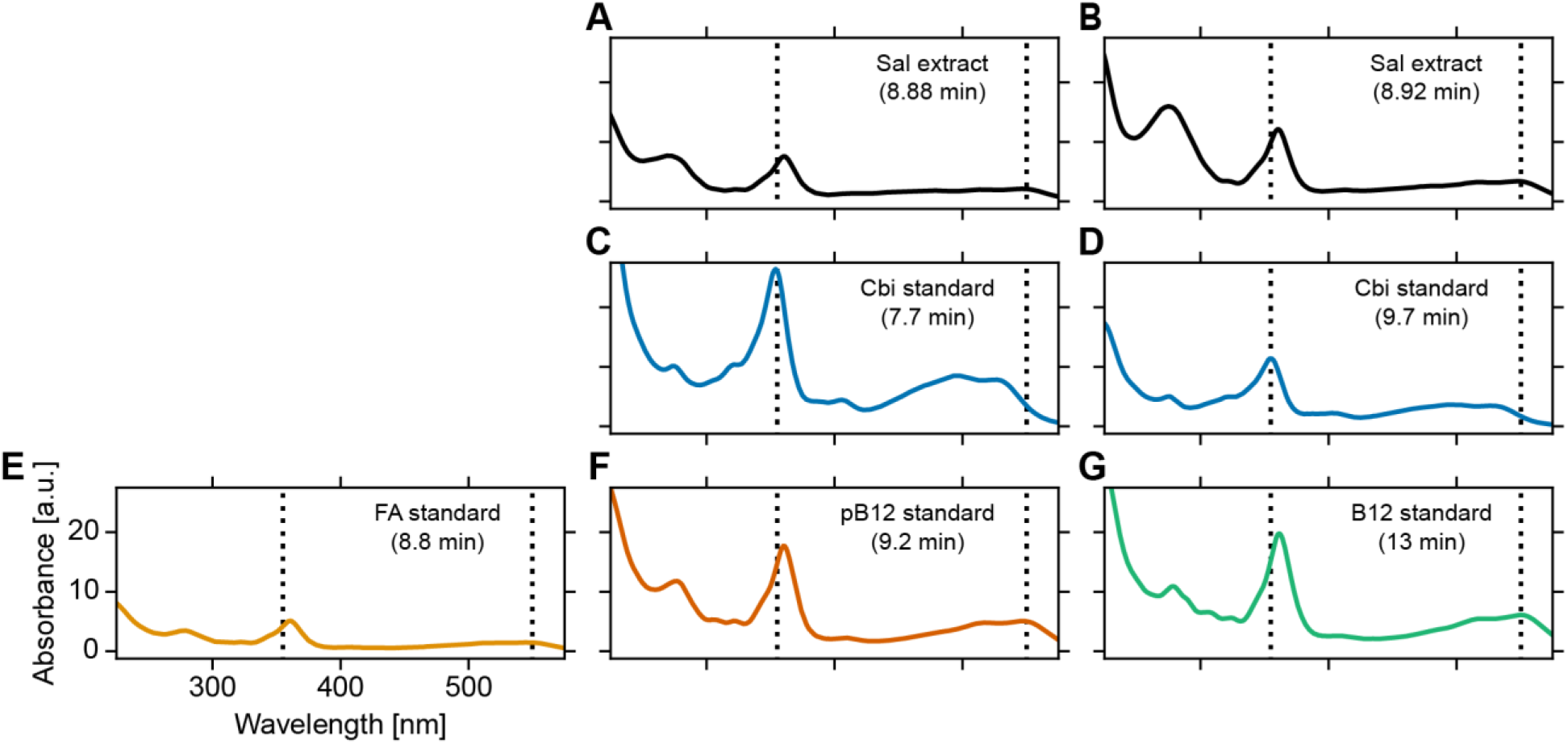
UV-Vis spectra of HPLC peaks Absorbance as a function of wavelength for selected HPLC peaks labeled in Fig. 3D. **A-B)** Peak spectra of Sal extracts. **C-G)** Peak spectra of known standards. Vertical dashed lines mark 355 nm and 550 nm, respectively.

**Supplementary Figure 5:**
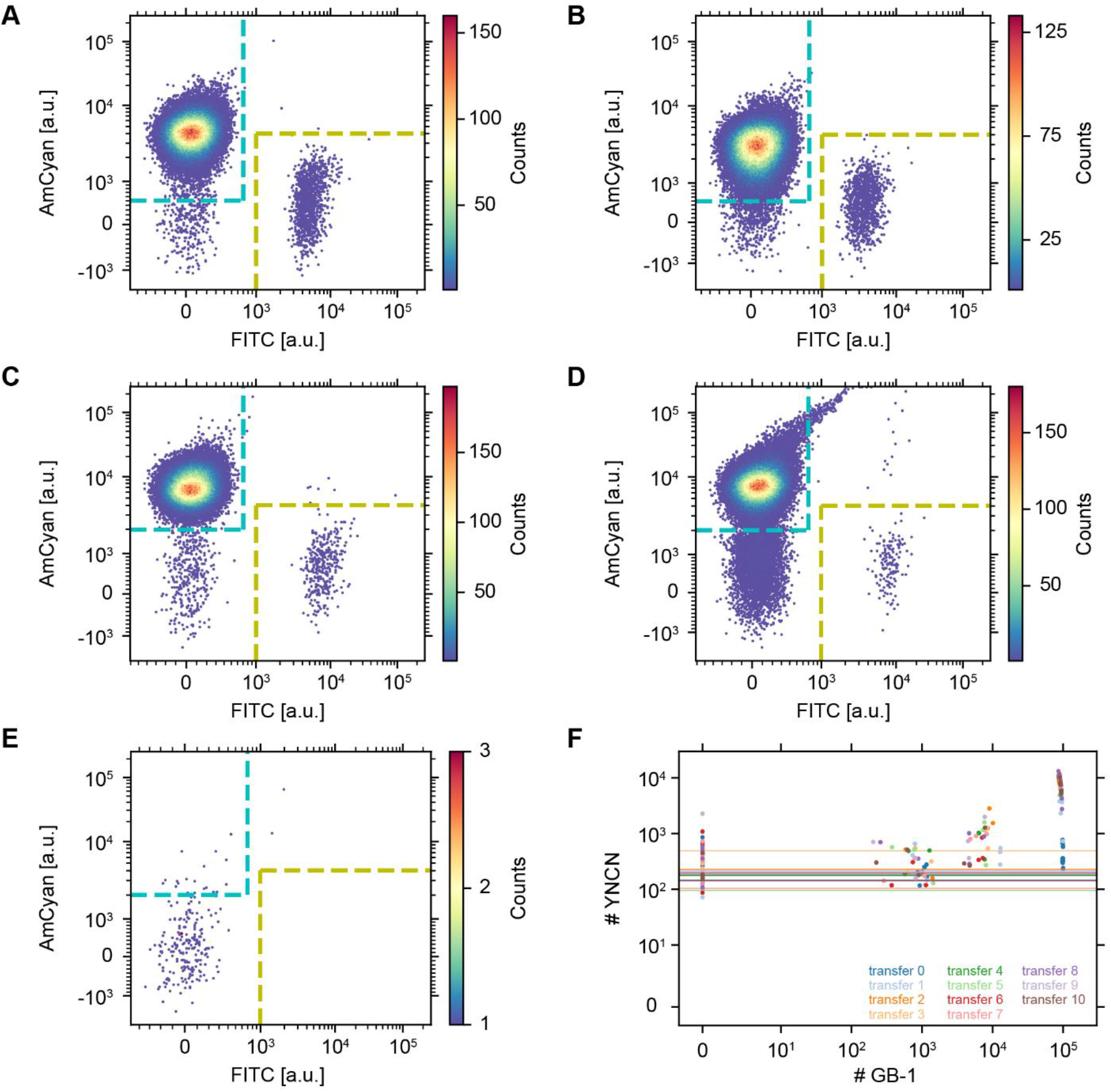
Classification of flow cytometry events **A-E)** AmCyan vs FITC scatter plots color-coded by event counts. AmCyan and FITC axes scales are logicle. Dashed lines mark gating conditions for YFP-negative-CFP-positive (cyan) and YFP-positive-CFP-negative (yellow) events, respectively. YFP-negative-CFP-negative events are located in the lower left quadrant of each plot, i.e., below the horizontal, dashed cyan lines and left of the vertical, dashed yellow lines. YFP-positive-CFP-positive events are located in the upper right quadrant of each plot, i.e., all events that do not fall into any of the other three gates. Notably, YFP-positive-CFP-positive events are very rare. Hence, no correction was applied, and YFP-positive-CFP-positive events were simply ignored for population ratio estimations. **A/C)** Mixed cultures of Sal/Dep at transfer 0 and transfer 1, respectively. **B/D)** Mixed cultures of Sal^GB-1^/Dep^GB-1^ at transfer 0 and transfer 1, respectively. A notable increase in YFP-negative-CFP-negative events can be seen at transfer 1 only if GB-1 cells are present in the mixed culture. **E)** Blank at transfer 1. **F)** Number of YFP-negative-CFP-negative (YNCN) events as function of the total number of GB-1 events detected in a culture. The total number of GB-1 events was estimated from YFP-negative-CFP-positive and YFP-positive-CFP-negative event counts and the known strain background identity (WT or GB-1). Solid lines mark YFP-negative-CFP-negative event counts of blanks. Both axes are linear below 10 and logarithmic thereafter. Cultures with a higher number of GB-1 events show a tendency to contain more YFP-negative-CFP-negative events (except for transfer 0, dark blue), suggesting that the occurrence of YFP-negative-CFP-negative events is mainly caused by strain background identity (WT or GB-1) rather than cobamide salvaging phenotype (salvager or dependent). No correction was applied, since adding the YFP-negative-CFP-negative events to the counts of GB-1 events only marginally altered the population ratio estimates.

**Supplementary Table 1:** Cobamide archetypes Counts of cobamide archetypes per phylum containing at least 10 genomes[30]. Same data (and same ordering, top-to-bottom in table corresponds to left-to-right in figure) as shown in Fig. 1B.

| Phylum | Producers | Salvagers | Dependents | Independents |
| --- | --- | --- | --- | --- |
| Nitrospinae | 0 | 2 | 8 | 0 |
| Candidatus Azambacteria | 0 | 0 | 10 | 0 |
| Candidatus Curtissbacteria | 0 | 0 | 9 | 1 |
| Candidatus Saccharibacteria | 0 | 0 | 2 | 8 |
| Chlorobi | 0 | 10 | 0 | 0 |
| Candidatus Sungbacteria | 0 | 0 | 6 | 5 |
| Ignavibacteriae | 0 | 0 | 12 | 0 |
| Candidatus Magasanikbacteria | 0 | 0 | 8 | 4 |
| Candidatus Staskawiczbacteria | 0 | 0 | 4 | 9 |
| Candidatus Taylorbacteria | 0 | 0 | 6 | 8 |
| Candidatus Gottesmanbacteria | 0 | 0 | 13 | 2 |
| Candidatus Omnitrophica | 0 | 2 | 14 | 0 |
| Candidatus Doudnabacteria | 0 | 0 | 15 | 1 |
| Candidatus Zambryskibacteria | 0 | 0 | 17 | 0 |
| Elusimicrobia | 1 | 3 | 12 | 1 |
| Candidatus Kaiserbacteria | 0 | 0 | 2 | 15 |
| Candidatus Uhrbacteria | 0 | 0 | 16 | 2 |
| Fibrobacteres | 0 | 1 | 18 | 0 |
| Candidatus Peregrinibacteria | 0 | 0 | 18 | 1 |
| Candidatus Daviesbacteria | 0 | 0 | 16 | 4 |
| Synergistetes | 6 | 9 | 5 | 0 |
| Aquificae | 0 | 0 | 13 | 8 |
| Candidatus Levybacteria | 0 | 0 | 17 | 5 |
| Nitrospirae | 8 | 7 | 6 | 1 |
| Candidatus Woesebacteria | 0 | 0 | 19 | 4 |
| Candidatus Giovannonibacteria | 0 | 0 | 19 | 8 |
| Candidatus Roizmanbacteria | 0 | 0 | 21 | 7 |
| Verrucomicrobia | 0 | 2 | 26 | 1 |
| Candidatus Yanofskybacteria | 0 | 0 | 34 | 0 |
| Thermotogae | 2 | 9 | 24 | 0 |
| Planctomycetes | 8 | 1 | 28 | 0 |
| Acidobacteria | 4 | 7 | 28 | 0 |
| Chlamydiae | 0 | 0 | 16 | 26 |
| Chloroflexi | 11 | 16 | 19 | 0 |
| Fusobacteria | 44 | 4 | 0 | 2 |
| Deinococcus-Thermus | 15 | 23 | 14 | 0 |
| Spirochaetes | 17 | 11 | 40 | 30 |
| Tenericutes | 0 | 1 | 26 | 115 |
| Cyanobacteria | 203 | 2 | 20 | 2 |
| Bacteroidetes | 6 | 324 | 689 | 21 |
| Actinobacteria | 1022 | 38 | 552 | 195 |
| Firmicutes | 646 | 473 | 744 | 275 |
| Proteobacteria | 2180 | 974 | 1423 | 258 |

**Supplementary Table 2:** Bacterial strains

| name | genotype | source |
| --- | --- | --- |
| Sal | *Escherichia coli* MG1655 ΔmetE | Kirmiz et al[52] |
| Dep | *Escherichia coli* MG1655 ΔmetE ΔcobUST ΔcobC | Mok et al[51] |
| Sal<sup>GB-1</sup> | *Escherichia coli* MG1655 GB-1 ΔmetE | This study. |
| Dep<sup>GB-1</sup> | *Escherichia coli* MG1655 GB-1 ΔmetE ΔcobUST ΔcobC | This study. |

**Supplementary Table 3:** Plasmids

| name | backbone | insert | source |
| --- | --- | --- | --- |
| pSG013 | Kan<sup>R</sup> marker, pBR322 origin, and rop gene | constitutive promoter (BBa_J23100, iGEM), RBS (BBa_B0034, iGEM), and mCerulean | This study. |
| pSG015 | Kan<sup>R</sup> marker, pBR322 origin, and rop gene | constitutive promoter (BBa_J23100, iGEM), RBS (BBa_B0034, iGEM), and mCitrine | This study. |
| pSG033 | Kan<sup>R</sup> marker, pBR322 origin, and rop gene | cobUST regulatory region (genomic region 500 bp upstream of cobUST start codon in *E. coli* MG1655 wt) and mCitrine | This study. |

## Notes

### Competing Interest Statement

The authors have declared no competing interest.

### Summary of Updates

Revised text and Fig. 1 to increase clarity.

## References

1. D’Souza, G., Shitut, S., Preussger, D., Yousif, G., Waschina, S., and Kost, C. (2018). Ecology and evolution of metabolic cross-feeding interactions in bacteria. Natural Product Reports 35, 455–488.

2. Gude, S., Pherribo, G.J., and Taga, M.E. (2020). Emergence of Metabolite Provisioning as a By-Product of Evolved Biological Functions. mSystems 5, e00259–00220.

3. Campbell, K., Herrera-Dominguez, L., Correia-Melo, C., Zelezniak, A., and Ralser, M. (2018). Biochemical principles enabling metabolic cooperativity and phenotypic heterogeneity at the single cell level. Current Opinion in Systems Biology 8, 97–108.

4. Fletcher, J.A., and Doebeli, M. (2009). A simple and general explanation for the evolution of altruism. Proceedings of the Royal Society B: Biological Sciences 276, 13–19.

5. Travisano, M., and Velicer, G.J. (2004). Strategies of microbial cheater control. Trends in Microbiology 12, 72–78.

6. Pande, S., Kaftan, F., Lang, S., Svatoš, A., Germerodt, S., and Kost, C. (2016). Privatization of cooperative benefits stabilizes mutualistic cross-feeding interactions in spatially structured environments. The ISME Journal 10, 1413–1423.

7. Smith, R.P., Doiron, A., Muzquiz, R., Fortoul, M.C., Haas, M., Abraham, T., Quinn, R.J., Barraza, I., Chowdhury, K., and Nemzer, L.R. (2019). The public and private benefit of an impure public good determines the sensitivity of bacteria to population collapse in a snowdrift game. Environmental Microbiology 21, 4330–4342.

8. Rankin, D.J., Bargum, K., and Kokko, H. (2007). The tragedy of the commons in evolutionary biology. Trends in Ecology & Evolution 22, 643–651.

9. D’Souza, G., Waschina, S., Pande, S., Bohl, K., Kaleta, C., and Kost, C. (2014). LESS IS MORE: SELECTIVE ADVANTAGES CAN EXPLAIN THE PREVALENT LOSS OF BIOSYNTHETIC GENES IN BACTERIA. Evolution 68, 2559–2570.

10. Freilich, S., Zarecki, R., Eilam, O., Segal, E.S., Henry, C.S., Kupiec, M., Gophna, U., Sharan, R., and Ruppin, E. (2011). Competitive and cooperative metabolic interactions in bacterial communities. Nature Communications 2, 589.

11. Harcombe, W. (2010). NOVEL COOPERATION EXPERIMENTALLY EVOLVED BETWEEN SPECIES. Evolution 64, 2166–2172.

12. Mee, M.T., Collins, J.J., Church, G.M., and Wang, H.H. (2014). Syntrophic exchange in synthetic microbial communities. Proceedings of the National Academy of Sciences 111, E2149–E2156.

13. Pacheco, A.R., Moel, M., and Segrè, D. (2019). Costless metabolic secretions as drivers of interspecies interactions in microbial ecosystems. Nature Communications 10, 103.

14. Morris, J.J., Lenski, R.E., and Zinser, E.R. (2012). The Black Queen Hypothesis: Evolution of Dependencies through Adaptive Gene Loss. mBio 3, e00036–00012.

15. Gore, J., Youk, H., and van Oudenaarden, A. (2009). Snowdrift game dynamics and facultative cheating in yeast. Nature 459, 253–256.

16. Gur, E., Biran, D., and Ron, E.Z. (2011). Regulated proteolysis in Gram-negative bacteria — how and when? Nature Reviews Microbiology 9, 839–848.

17. Wolfe, A.J. (2005). The Acetate Switch. Microbiology and Molecular Biology Reviews 69, 12–50.

18. Villalba, G., Segarra, M., Fernández, A.I., Chimenos, J.M., and Espiell, F. (2002). A proposal for quantifying the recyclability of materials. Resources, Conservation and Recycling 37, 39–53.

19. Neilands, J.B. (1995). Siderophores: Structure and Function of Microbial Iron Transport Compounds (*). Journal of Biological Chemistry 270, 26723–26726.

20. Butaitė, E., Baumgartner, M., Wyder, S., and Kümmerli, R. (2017). Siderophore cheating and cheating resistance shape competition for iron in soil and freshwater Pseudomonas communities. Nature Communications 8, 414.

21. Kramer, J., Özkaya, Ö., and Kümmerli, R. (2020). Bacterial siderophores in community and host interactions. Nature Reviews Microbiology 18, 152–163.

22. Johnson, M.D.L., Echlin, H., Dao, T.H., and Rosch, J.W. (2015). Characterization of NAD salvage pathways and their role in virulence in Streptococcus pneumoniae. Microbiology 161, 2127–2136.

23. Melnick, J., Lis, E., Park, J.-H., Kinsland, C., Mori, H., Baba, T., Perkins, J., Schyns, G., Vassieva, O., Osterman, A., et al. (2004). Identification of the Two Missing Bacterial Genes Involved in Thiamine Salvage: Thiamine Pyrophosphokinase and Thiamine Kinase. J Bacteriol 186, 3660–3662.

24. Nygaard, P. (1993). Purine and Pyrimidine Salvage Pathways. In Bacillus subtilis and Other Gram-Positive Bacteria. (American Society of Microbiology).

25. Sekowska, A., Dénervaud, V., Ashida, H., Michoud, K., Haas, D., Yokota, A., and Danchin, A. (2004). Bacterial variations on the methionine salvage pathway. BMC Microbiology 4, 9.

26. Yuan, Y., Zallot, R., Grove, T.L., Payan, D.J., Martin-Verstraete, I., Šepić, S., Balamkundu, S., Neelakandan, R., Gadi, V.K., Liu, C.-F., et al. (2019). Discovery of novel bacterial queuine salvage enzymes and pathways in human pathogens. Proceedings of the National Academy of Sciences 116, 19126–19135.

27. Datta, M.S., Sliwerska, E., Gore, J., Polz, M.F., and Cordero, O.X. (2016). Microbial interactions lead to rapid micro-scale successions on model marine particles. Nature Communications 7, 11965.

28. Escalante-Semerena, J.C. (2007). Conversion of Cobinamide into Adenosylcobamide in Bacteria and Archaea. J Bacteriol 189, 4555–4560.

29. Contreras, H., Chim, N., Credali, A., and Goulding, C.W. (2014). Heme uptake in bacterial pathogens. Current Opinion in Chemical Biology 19, 34–41.

30. Shelton, A.N., Seth, E.C., Mok, K.C., Han, A.W., Jackson, S.N., Haft, D.R., and Taga, M.E. (2019). Uneven distribution of cobamide biosynthesis and dependence in bacteria predicted by comparative genomics. The ISME Journal 13, 789–804.

31. Pande, S., Merker, H., Bohl, K., Reichelt, M., Schuster, S., de Figueiredo, L.F., Kaleta, C., and Kost, C. (2014). Fitness and stability of obligate cross-feeding interactions that emerge upon gene loss in bacteria. The ISME Journal 8, 953–962.

32. Noto Guillen, M., Rosener, B., Sayin, S., and Mitchell, A. (2021). Assembling stable syntrophic Escherichia coli communities by comprehensively identifying beneficiaries of secreted goods. Cell Systems.

33. Anderson, P.J., Lango, J., Carkeet, C., Britten, A., Kräutler, B., Hammock, B.D., and Roth, J.R. (2008). One Pathway Can Incorporate either Adenine or Dimethylbenzimidazole as an α-Axial Ligand of B12 Cofactors in Salmonella enterica. J Bacteriol 190, 1160–1171.

34. Chan, C.H., and Escalante-Semerena, J.C. (2011). ArsAB, a novel enzyme from Sporomusa ovata activates phenolic bases for adenosylcobamide biosynthesis. Molecular Microbiology 81, 952–967.

35. Degnan, Patrick H., Barry, Natasha A., Mok, Kenny C., Taga, Michiko E., and Goodman, Andrew L. (2014). Human Gut Microbes Use Multiple Transporters to Distinguish Vitamin B12 Analogs and Compete in the Gut. Cell Host & Microbe 15, 47–57.

36. Crofts, Terence S., Seth, Erica C., Hazra, Amrita B., and Taga, Michiko E. (2013). Cobamide Structure Depends on Both Lower Ligand Availability and CobT Substrate Specificity. Chemistry & Biology 20, 1265–1274.

37. Hazra, Amrita B., Tran, Jennifer L.A., Crofts, Terence S., and Taga, Michiko E. (2013). Analysis of Substrate Specificity in CobT Homologs Reveals Widespread Preference for DMB, the Lower Axial Ligand of Vitamin B12. Chemistry & Biology 20, 1275–1285.

38. Hazra, A.B., Han, A.W., Mehta, A.P., Mok, K.C., Osadchiy, V., Begley, T.P., and Taga, M.E. (2015). Anaerobic biosynthesis of the lower ligand of vitamin B_12_. Proceedings of the National Academy of Sciences 112, 10792–10797.

39. Davies, J. (1994). Inactivation of Antibiotics and the Dissemination of Resistance Genes. Science 264, 375–382.

40. Jones, C.M., Hernández Lozada, N.J., and Pfleger, B.F. (2015). Efflux systems in bacteria and their metabolic engineering applications. Applied Microbiology and Biotechnology 99, 9381–9393.

41. Cowan, S.E., Gilbert, E., Liepmann, D., and Keasling, J.D. (2000). Commensal Interactions in a Dual-Species Biofilm Exposed to Mixed Organic Compounds. Applied and Environmental Microbiology 66, 4481–4485.

42. Smith, P., and Schuster, M. (2019). Public goods and cheating in microbes. Current Biology 29, R442–R447.

43. Herring, C.D., Raghunathan, A., Honisch, C., Patel, T., Applebee, M.K., Joyce, A.R., Albert, T.J., Blattner, F.R., van den Boom, D., Cantor, C.R., et al. (2006). Comparative genome sequencing of Escherichia coli allows observation of bacterial evolution on a laboratory timescale. Nature Genetics 38, 1406–1412.

44. Cheng, K.-K., Lee, B.-S., Masuda, T., Ito, T., Ikeda, K., Hirayama, A., Deng, L., Dong, J., Shimizu, K., Soga, T., et al. (2014). Global metabolic network reorganization by adaptive mutations allows fast growth of Escherichia coli on glycerol. Nature Communications 5, 3233.

45. Burkovski, A., and Krämer, R. (2002). Bacterial amino acid transport proteins: occurrence, functions, and significance for biotechnological applications. Applied Microbiology and Biotechnology 58, 265–274.

46. Jojima, T., Omumasaba, C.A., Inui, M., and Yukawa, H. (2010). Sugar transporters in efficient utilization of mixed sugar substrates: current knowledge and outlook. Applied Microbiology and Biotechnology 85, 471–480.

47. Yi, S., Seth, E.C., Men, Y.-J., Stabler, S.P., Allen, R.H., Alvarez-Cohen, L., and Taga, M.E. (2012). Versatility in Corrinoid Salvaging and Remodeling Pathways Supports Corrinoid-Dependent Metabolism in Dehalococcoides mccartyi. Applied and Environmental Microbiology 78, 7745–7752.

48. Estrela, S., Morris, J.J., and Kerr, B. (2016). Private benefits and metabolic conflicts shape the emergence of microbial interdependencies. Environmental Microbiology 18, 1415–1427.

49. Lindsay, R.J., Pawlowska, B.J., and Gudelj, I. (2019). Privatization of public goods can cause population decline. Nature Ecology & Evolution 3, 1206–1216.

50. Gray, M.J., and Escalante-Semerena, J.C. (2009). The cobinamide amidohydrolase (cobyric acid-forming) CbiZ enzyme: a critical activity of the cobamide remodelling system of Rhodobacter sphaeroides. Molecular Microbiology 74, 1198–1210.

51. Mok, K.C., Sokolovskaya, O.M., Nicolas, A.M., Hallberg, Z.F., Deutschbauer, A., Carlson, H.K., Taga, M.E., and Lemon, K.P. (2020). Identification of a Novel Cobamide Remodeling Enzyme in the Beneficial Human Gut Bacterium Akkermansia muciniphila. mBio 11, e02507–02520.

52. Kirmiz, N., Galindo, K., Cross, K.L., Luna, E., Rhoades, N., Podar, M., Flores, G.E., and Liu, S.-J. (2020). Comparative Genomics Guides Elucidation of Vitamin B_12_ Biosynthesis in Novel Human-Associated *Akkermansia* Strains. Applied and Environmental Microbiology 86, e02117–02119.

53. Baba, T., Ara, T., Hasegawa, M., Takai, Y., Okumura, Y., Baba, M., Datsenko, K.A., Tomita, M., Wanner, B.L., and Mori, H. (2006). Construction of Escherichia coli K-12 in-frame, single-gene knockout mutants: the Keio collection. Molecular Systems Biology 2, 2006.0008.

54. Datsenko, K.A., and Wanner, B.L. (2000). One-step inactivation of chromosomal genes in Escherichia coli K-12 using PCR products. Proceedings of the National Academy of Sciences 97, 6640–6645.

55. Gibson, D.G., Young, L., Chuang, R.-Y., Venter, J.C., Hutchison, C.A., and Smith, H.O. (2009). Enzymatic assembly of DNA molecules up to several hundred kilobases. Nature Methods 6, 343–345.

56. (2019). Python Software Foundation, Python v.3.8.

57. Harris, C.R., Millman, K.J., van der Walt, S.J., Gommers, R., Virtanen, P., Cournapeau, D., Wieser, E., Taylor, J., Berg, S., Smith, N.J., et al. (2020). Array programming with NumPy. Nature 585, 357–362.

58. Virtanen, P., Gommers, R., Oliphant, T.E., Haberland, M., Reddy, T., Cournapeau, D., Burovski, E., Peterson, P., Weckesser, W., Bright, J., et al. (2020). SciPy 1.0: fundamental algorithms for scientific computing in Python. Nature Methods 17, 261–272.

59. Castillo-Hair, S.M., Sexton, J.T., Landry, B.P., Olson, E.J., Igoshin, O.A., and Tabor, J.J. (2016). FlowCal: A User-Friendly, Open Source Software Tool for Automatically Converting Flow Cytometry Data from Arbitrary to Calibrated Units. ACS Synthetic Biology 5, 774–780.

60. Sokolovskaya, O.M., Mok, K.C., Park, J.D., Tran, J.L.A., Quanstrom, K.A., Taga, M.E., and Ribbe, M.W. (2019). Cofactor Selectivity in Methylmalonyl Coenzyme A Mutase, a Model Cobamide-Dependent Enzyme. mBio 10, e01303–01319.

